# Addressing the Embeddability Problem in Transition Rate Estimation

**DOI:** 10.1101/707919

**Authors:** Curtis Goolsby, James Losey, Yuchen Xu, Marie-Christine Düker, Mila Getmansky Sherman, David S. Matteson, Mahmoud Moradi

## Abstract

Markov State Models (MSM) and related techniques have gained significant traction as a tool for analyzing and guiding molecular dynamics (MD) simulations due to their ability to extract structural, thermodynamic, and kinetic information on proteins using computationally feasible MD simulations. The MSM analysis often relies on spectral decomposition of empirically generated transition matrices. Here, we discuss an alternative approach for extracting the thermodynamic and kinetic information from the so-called rate/generator matrix rather than the transition matrix. Although the rate matrix itself is built from the empirical transition matrix, it provides an alternative approach for estimating both thermodynamic and kinetic quantities, particularly in diffusive processes. We particularly discuss a fundamental issue with this approach, known as the embeddability problem and offer ways to address this issue. We describe eight different methods to overcome the embeddability problem, including a novel approach developed for this work. The algorithms were tested on data from a one-dimensional toy model to show the workings of these methods and discuss the robustness of each method in terms of its dependence in lag time and trajectory length.

## 1. INTRODUCTION

Proteins and other biological macromolecules are associated with complex conformational spaces that are virtually impossible to be fully characterized at the atomic level using current experimental and computational tools [1]. With increasing computational capabilities due to hardware improvements, all-atom molecular dynamics simulations are used to explore the possibility of investigating the important regions of free energy landscapes of proteins and other biomolecules[2 − −4]. The tools provided by computational methods are limited by their computational costs. With this limitation, it is of significant interest to employ statistical techniques, which allow the inference of relevant thermodynamic and kinetic properties from shorter, less costly simulations. Markov State Models or MSMs[5 − −10] provide some of the most powerful tools for analyzing the ensembles of short molecular dynamics trajectories to extract information on both thermodynamics and kinetics of complex biomolecular systems and predict the behavior of such systems at much longer timescales than would be possible to simulate with current computing capabilities. These methods, however, are based on assumptions and simplifications that introduce limitations to the reliability and interpretability of these methods[11 − −13].

Building MSMs involves (1) discretization of conformational space, (2) extracting transition statistics from simulation trajectories, and (3) analyzing the transition statistics to estimate kinetic and thermodynamic properties[14]. Here, we only focus on the third component, which assumes an empirical transition matrix has been generated. Here we discuss a known[15] but less commonly used approach for analyzing empirical transition matrices that in some cases could provide an alternative approach to the more common eigendecomposition technique. The latter relies on the eigenvectors and eigenvalues of the empirical transition matrix to estimate thermodynamic and kinetic properties[16]. The approach we will discuss here, however, is based on building a rate matrix to estimate the kinetic and thermodynamic quantities using standard methods from chemical kinetics literature (Figure 1). In the finance literature, the rate matrix is known as the generator matrix[17]. Producing a rate matrix from the empirical transition matrix is known to be associated with the embeddability problem in the field of finance. The embeddability problem is found in taking the matrix logarithm of a time-dependent transition probability matrix in order to determine the generator matrix or the rate matrix, as known in chemical kinetics literature. In theory, the lag-time dependent transition probability matrix should only have positive eigenvalues; however, in practice, insufficient sampling can result in the presence of negative eigenvalues. Thus, its matrix logarithm has non-real values and is an invalid generator matrix. Solving this embeddability problem involves accurately predicting the true generator matrix from an invalid generator matrix. We explored the validity of various algorithms in the chemical literature as well as from the field of finance. These methods were applied to data generated from a simple one-dimensional toy model, and compared to assess the relative performance of proposed algorithms empirically.

**FIG. 1.**
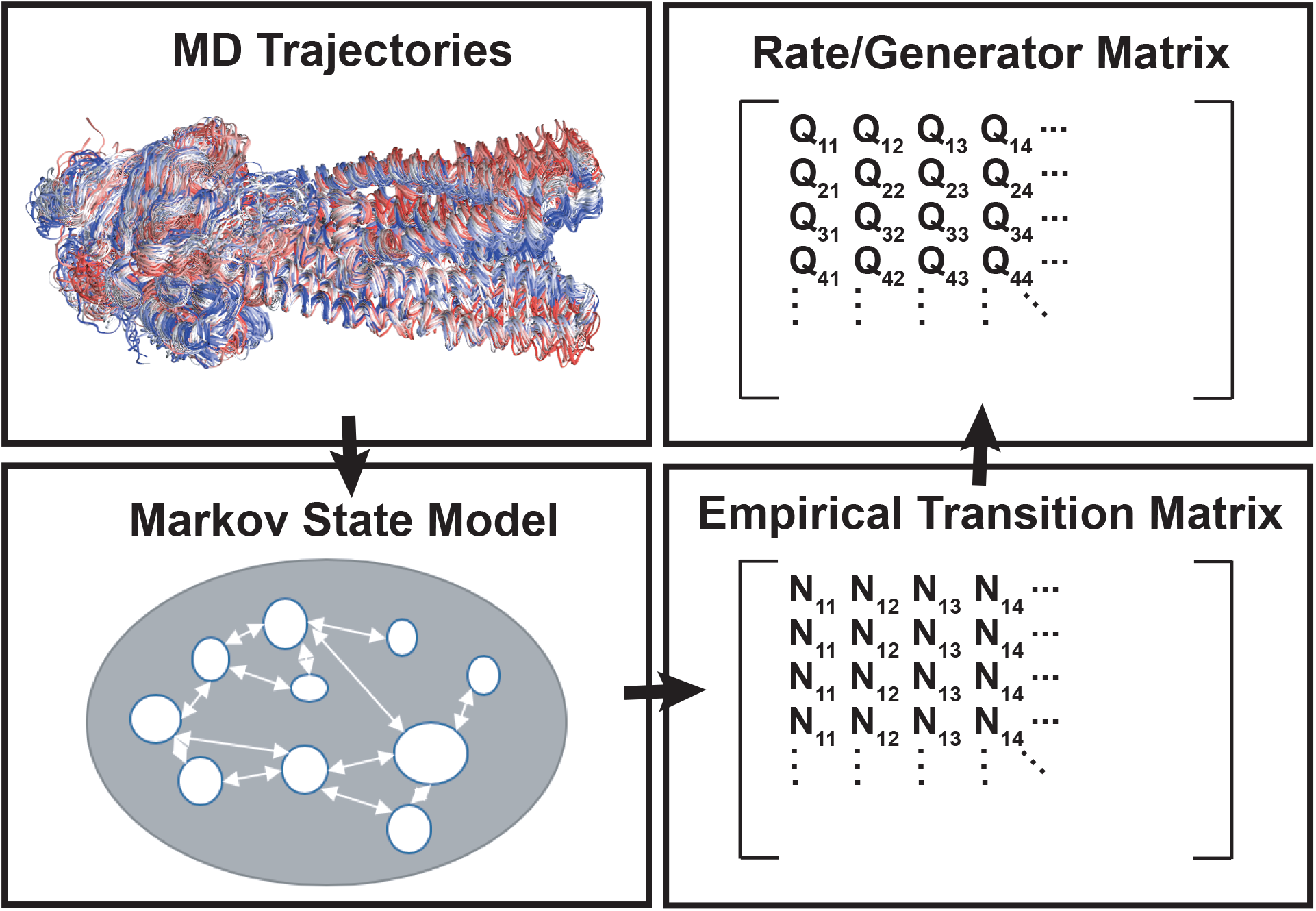
The schematic representation of the proposed approach to analyze the MSM data. MD trajectories are first used within a clustering scheme to generate a Markov state model and build an empirical transition matrix using a given lag time (see MSM literature[7]). Instead of spectral decomposition of the transition matrix, however, we propose to estimate a lag-time independent rate/generator matrix from the empirical transition matrix obtained from the MD trajectories.

## II. BACKGROUND

Enhanced sampling[18] has found success in the field of computational chemistry/biophysics as an effective remedy for the costs involved with simulating large molecular systems. Methods such as umbrella sampling (US)[19 − −22], metadynamics[23 − −26], and replica exchange[27 − −31] have grown increasingly popular by aiming to enhance the sampling of configuration space by manipulating the energetics of the system. Enhancing the sampling by biasing the potential or kinetic energy, of course requires post-analysis reweighting techniques such as weighted histogram analysis method or multistate Bennett acceptance ratio method to determine the thermodynamic and kinetic quantities such as free energy and mean first passage time (MFPT). However, it is reasonable to assume that the reweighting schemes cannot fully remove the inherent bias in the MD data generated using enhanced sampling techniques. As a consequence, an alternative approach to characterize the thermodynamic and kinetic properties of biomolecular systems has been suggested that relies on Markovian analysis of transition probabilities between discrete states obtained from short but numerous unbiased MD trajectories. These methods are often known as Markov State Models or MSMs[5 − −10].

In this paper we explore the possibility of different algorithmic techniques for solving what is known as the “embeddability problem”[32, 33] in financial literature. We present a comparison between eight algorithms, used for predicting bond rating transitions, implemented in Inamura[34] and an extension by Marada[35] of Inamura’s work with known and largely used methods of estimating free energies, as well as a maximum likelihood approach based on Hummer *et al*.[15]. In order to explore the efficacy of each of these methods we used a 1D bistable potential model. It is important to stress the comparison between algorithms found in finance for performing work on bond rating matrices, and our work here. The underlying mathematics is identical, and in fact there may be hidden gems in other seemingly unrelated fields for the application of this work.

Determination of kinetic properties is fundamental to the understanding of molecular systems. In order to use molecular dynamics as more than a virtual microscope, detailed calculations are necessary in order to search the conformational space for properties and conformations of interest. Various methods, such as those mentioned above, have been developed to improve the sampling in order to determine thermodynamic and kinetic properties of systems. These models are largely based upon defining a collective variable space[36, 37] in which to reduce the dimensionality of the sampling space. The methods then address how to sample more efficiently along a predefined collective variable.

MSMs[11, 38–40] allow for the reduction of the complexities of a dynamic molecular system into a lower dimensional model. The conformational space can be reduced by clustering and then brought into a square empirical transition matrix allowing for the determination of various parameters of interest such as the relative free energies and diffusion constants among others. MSMs as with other computational methodologies still require a sufficient amount of sampling in order to obtain the parameters of interest. The present work adapts several algorithms from the field of finance in order to compare their efficacy with that of the state of the art in the chemical physics literature for addressing problems arising from insufficient sampling. In finance, MSMs are built in order to determine the probabilities of transition between rating grades for bonds[34]. Often there is insufficient data in the bond marketplace to observe transitions from every rating to every other rating in the same fashion that the conformational space of a protein is too complex in order to sample every transition with MD[34]. The discrete nature of the problem and the assumption of memoryless-ness cause the same models to work in both finance and chemical physics.

The algorithms, described below, are applied to a 1D bistable toy model. We test the efficacy of the various algorithms under different conditions, first in sufficient sampling to show that the algorithms do not distort the correct answer, and then in the case of increasing lag times as well as the case of insufficient data points. By doing this we are able to show the various strengths and weaknesses of each algorithm and hope to provide guidance for future researchers in their decision making about this matter.

Let us assume that we can assign any given sampled conformation from a set of MD simulations to one of the *K* well-defined states that are determined by clustering or other dimensionality reduction techniques. The empirical transition matrix, *N*, is defined as a *K* × *K* matrix, where *N*_*i, j*_ is the number of observations of system being at state *i* at time *t* and being at state *j* at time *t* + *τ*, in which *τ* is a given lag time. An estimator **P** for the transition matrix can be obtained by normalizing *N*.

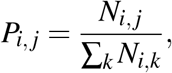

where *P*_*i, j*_ is the empirical probability of observing the system at state *j* at time *t* + *τ*, given the system is observed at state *i* at time *t*.

From the transition matrix **P**, we can find the rate/generator matrix **Q**. While the **P** matrix is a more common quantity to work with in Markov chain based models such as MSM, the rate matrix is more well-known in chemical kinetics literature and is the focus of our work here. **Q** is relevant for continuous time models. There are advantages in using **Q** over **P** for such models; e.g., **P** is by construct dependent on the lag time, while **Q** is not.

Transition matrix **P** is a function of *τ* and is related to rate/generator matrix **Q** by **P**(*τ*) = exp(**Q***τ*). The matrix **Q** can thus be estimated from **P** within the Markovian approximation using the relation

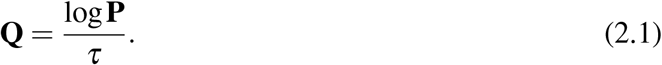

However, the elements of the matrix **Q** need to be real-valued and satisfy the criteria

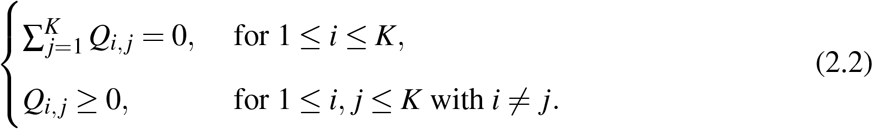

to be a valid generator matrix. The criteria in (2.2) lead to

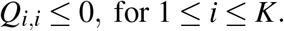

Additional criteria need to be imposed in order to guarantee the validity of detailed balance relation, which is a common feature of equilibrium processes. Put simply, at equilibrium each process should be equilibrated by its reverse process. The detailed balance or reversibility feature then allows for the definition of free energies based on the equilibrium probability of states

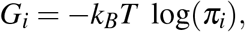

where *k*_*B*_ is the Boltzmann constant and *T* is the temperature, *G*_*i*_ is the free energy of state *i* and *π*_*i*_ is the probability of observing the system at state *i* at equilibrium. The free energy between any two states *i* and *j*, Δ*G*_*i, j*_ = *G*(*j*) − *G*(*i*), can be calculated from **Q** using the detailed balance relation:

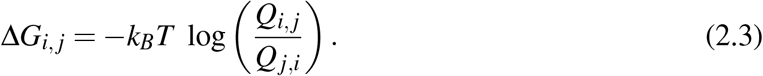

Another interesting feature of relevance to molecular processes is the diffusivity, where no jump is allowed beyond the immediate neighbors of a state. The diffusivity condition is satisfied when the generator matrix **Q** is tridiagonal. For a diffusive process, the generator matrix can be used to determine the diffusion constant as[15]

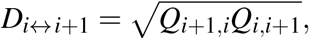

where *D*_*i*↔*i*+1_ is related to the continuous diffusion constant of a diffusive process by the following approximate relationship:

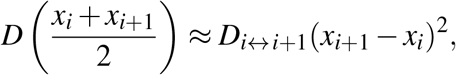

where *x*_*i*_ represents the position along “reaction coordinate” associated with state *i*. If the states are equidistantly distributed along the reaction coordinate, we can estimate

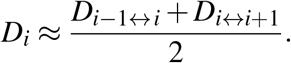

which leads to *D*(*x*) = *D*_*i*_Δ*x*^2^, in which Δ*x* = *x*_*i*_ − *x*_*i*−1_ for all *i*.

The diffusion constant and free energy of states can be used to determine the mean first passage time (MFPT). This is the time of transition from the reactant (i.e, free energy minimum at *x*_*R*_) to the product (i.e., free energy minimum at *x*_*P*_, with an effective infinite free energy at *x*_0_). Lifson and Jackson[41] and others[42] have shown the MFPT can be calculated using

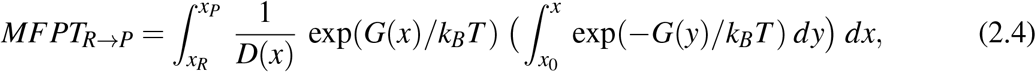

where the free energy and diffusion constant along *x* are described by *G*(*x*) and *D*(*x*), respectively.

The MFPT can be estimated from *G*_*i*_ and *D*_*i*_ values as

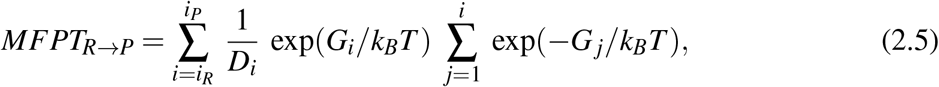

where *i*_*P*_ and *i*_*R*_ are the reactant and product states.

## III. THE EMBEDDABILITY PROBLEM

For a given empirical transition matrix, it has been shown that with a sufficient but not necessary condition[43], an exact generator matrix exists such that 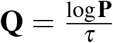 as in (2.1). However, the resulting **Q** may not be a valid generator matrix. For instance, consider states *i* and *j* such that transitions between the two are possible given infinite time but there is no recorded transition from *i* to *j* in the empirical transition matrix. Unfortunately, this is not an uncommon case for empirical transition matrices, due to under sampling certain transitions. Such behavior in empirical transition matrices would lead to empirical transition matrices that do not not satisfy the criteria in (2.2). One may even calculate negative eigenvalues for the empirical transition matrix, which would result in a non-real generator matrix.

In Section IV below, we present several methods to address this problem, namely either adjusting directly the generator matrix produced by the matrix logarithm of our insufficiently sampled transition matrix, performing a maximum likelihood estimation, or using a Markov chain Monte Carlo technique.

## IV. ALGORITHMS

We present here a wide range of algorithms whose comparison can be found in Section VI below. These algorithms range from very simple methods such as the Diagonal Adjustment algorithm to more intricate methods such as Expectation Maximization. Besides using existing algorithms suggested in related literature for comparison, we also introduce a novel algorithm in order to estimate the generator matrix **Q**. The procedure is called Polynomial Adjustment (PA) algorithm and is based on the common eigenvector structure of **Q** and the transition matrix **P**.

### A. Diagonal Adjustment (DA)

The Diagonal Adjustment (DA)[34] algorithm is a simple and sometimes effective solution to the embeddability problem. The algorithm adjusts the generator matrix in two steps:

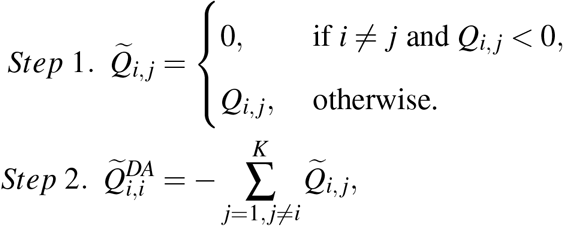

where 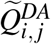 denotes the elements of an estimated generator matrix obtained by DA algorithm.

### B. Weighted Adjustment (WA)

The Weighted Adjustment (WA)[34] is another simple algorithm very similar to the DA. The algorithm adjusts the generator matrix in two steps.

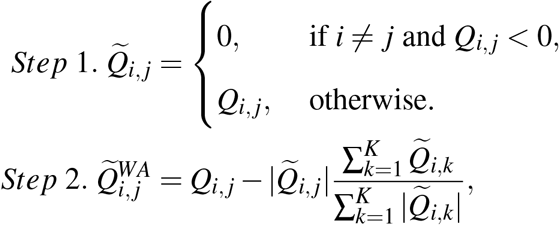

where 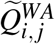 denotes the elements of an estimated generator matrix obtained by WA algorithm.

### C. Quasi-Optimization of the Generator (QOG)

DA and WA are very similar methodologies. Unfortunately, they are not based upon an optimization strategy and become increasingly hard to trust in sparse data situations due to their unphysicality. To this end, Krenin and Sidelnikova[44] have extended the above work by implementing a post-adjustment optimization method called quasi-optimization of the generator (QOG). QOG works by first noting that the generator has a restriction on each row which allows the problem to be split into *K* distinct minimization problems of the sum of the squared deviation between log **P** and **Q***τ*. Thus we can write the problem as

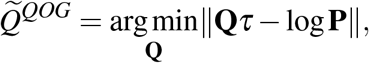

where ∥ · ∥denotes the so-called Frobenius norm defined as 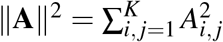 for a real-valued matrix **A** = (*A*_*i, j*_)_*i, j*=1,…,*K*_. By reducing the problem to a distinct problem for each row, we can define

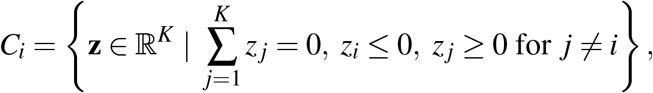

where **z** represents possible valid values for the *i*’th row of **Q**. The optimum **z** can be determined from the *i*’th row of log(**P**) (denoted by **a**) as

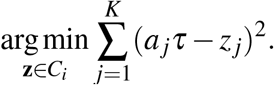

An easy implementation of this algorithm can be found in the Jupyter notebook contained in the supplementary information.

### D. Component-Wise Optimization (CWO)

Component-Wise Optimization (CWO)[35] is a somewhat more complex version of DA or WA which bears resemblance to QOG. The general idea is to divide the problem into K-1 separate optimization problems. First, a DA or WA is performed followed by optimizing each individual value in the generator according to

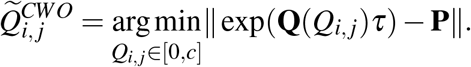

This is followed by rebalancing the row to maintain a valid generator matrix. Constant *c* determines the desired convergence; the smaller the *c*, the faster the convergence defined as 0.0001 in our work. Marada goes on to note that the CWO is not capable of distinguishing between local and global minima and as such should be used judiciously, possibly as a way to further optimize results from other algorithms[35].

### E. Expectation Maximization (EM)

The Expectation Maximization (EM)[34] algorithm is based on iterating two steps, the Expectation-step and the Maximization-step. To be more precise, recall that *N*_*i, j*_ denotes the number of transitions from state *i* to *j*. We further write *R*_*i*_ for the number of time steps the system stays in state *i*.

Then, write

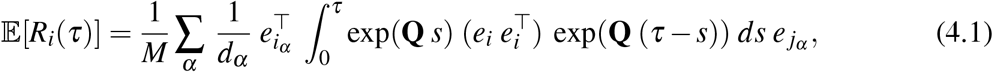

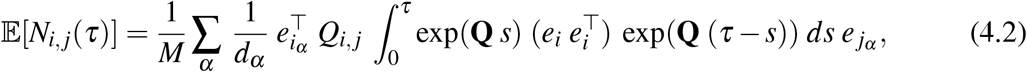

where *α* is an index pointing to a specific transition from state *i*_*α*_ (observed at time *t*_*α*_) to state *j*_*α*_ (observed at time *t*_*α*_ + *τ*), and *M* is the total number of transition observations for a given lag time *τ*. Furthermore, *e*_*i*_ denotes a unit vector with the *i*’th element being one and the rest of elements being zero, we write *e*^┬^ for the transpose of *e*. The quantity *d*_*α*_ is defined by

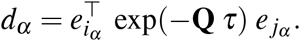

The relations (4.1) and (4.2) allow to estimate the expected values of *N*_*i, j*_ and *R*_*i*_ as a function of the generator matrix **Q**. The generator matrix is then estimated as

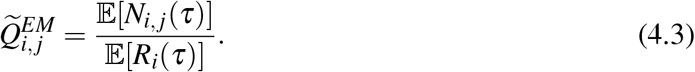

The procedure of the EM is done by calculating (4.1) and (4.2) for each element, and then using (4.3) to construct a new generator matrix. The iteration proceeds until convergence completes the algorithm.

### F. Maximum Likelihood Estimator (MLE)

A common optimization approach is to use maximum likelihood estimation (MLE), in this case to determine a generator matrix to maximize the likelihood of observing all of the transitions, *α* (summarized in the empirical transition matrix for a given lag time *τ*)[15]. The relation,

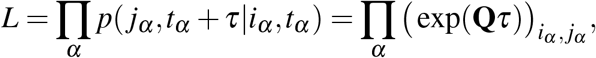

which leads to

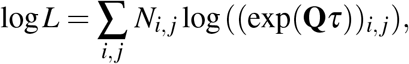

in which the off-diagonal elements of **Q** matrix are varied but stay positive by construct and the diagonal elements are determined by

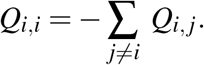

Other criteria can be easily imposed on **Q** such as reversibility (to satisfy the detailed balance) and diffusivity. To satisfy both reversibility and diffusivity, for instance, only *Q*_*i,i*±1_ elements are varied and the other elements are determined by

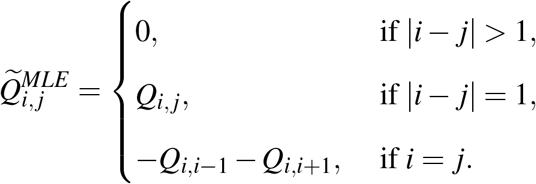

### G. Quadratic Programming (QP)

A quadratic programming approach was proposed by D.T. Crommelin and E. Vanden-Eijnden[45] to take into account the eigen-structures of the transition probability matrix **P** and the generator matrix **Q**. Indeed, if **P** has the eigendecomposition

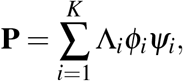

where **P*φ***_*i*_ = Λ_*i*_*φ*_*i*_, *Ψ*_*i*_**P** = Λ_*i*_*Ψ*_*i*_, Λ_*i*_ ∈ ℂ, ∀*i* = 1, …, *K*, then with *λ*_*i*_ = *τ*^−1^ log(Λ_*i*_),

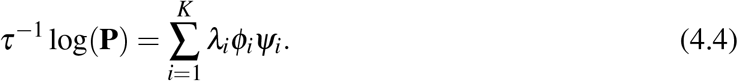

Note that (*φ*_*i*_, *Ψ*_*i*_, Λ_*i*_, *λ*_*i*_), for 1 ≤ *i* ≤ *K*, can be complex-valued.

Since the generator matrix **Q** has a similar eigen-structure as *τ*^−1^ log(**P**) given in (4.4), an estimator 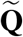 can be obtained by solving the following optimization problem

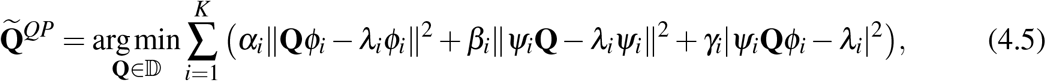

where *α*_*i*_, *β*_*i*_, *γ*_*i*_ for *i* = 1, …, *K* are positive weights chosen to stabilize the algorithm numerically, and domain 𝔻 is a subspace of ℝ ^*K*×*K*^ defined by (2.2). The norm-operators and the absolute values in (4.5) are compatible with complex cases. Note that the objective function in (4.5) is a quadratic function of the entries **Q**_*i, j*_ for all 1 ≤ *i, j* ≤ *K*, and (2.2) only imposes linear constraints to the domain space D, the problem (4.5) can thus be well handled by quadratic programming[46].

### H. Polynomial Adjustment (PA)

We propose here a novel routine of polynomial adjustment (PA) that not only meets the generator matrix constraints in (2.2) but also strictly maintains the eigenvectors structure.

The generator matrix **Q** can be expressed as a polynomial of the transition probability matrix **P** with order at most *K* − 1, or equivalently, the vectorization of the generator matrix can be given by a linear combination of the column vectors of the matrix

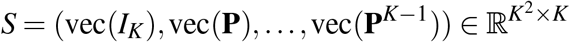

with vec(·) the operator that stacks all the columns of a matrix, and *I*_*K*_ the *K* × *K* identity matrix. It is then possible to parametrize the adjustment 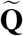 as

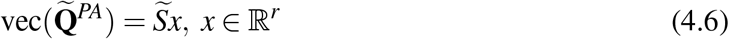

with 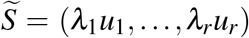 the column space basis of *S*. The singular value decomposition of *S* yields non-increasing singular values *λ*_1_, …, *λ*_*K*_ and corresponding left singular eigenvectors 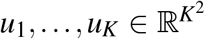, while *r* is the index that satisfies *λ*_*r*_ *> ε* ≥ *λ*_*r* + 1_ (set *λ* + *K*+1 := 0) for a given threshold *ε* ≥ 0 for numeric stability. With **Q** based on the power series expression of matrix logarithm of **P**, the minimizer 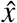 in the optimization problem

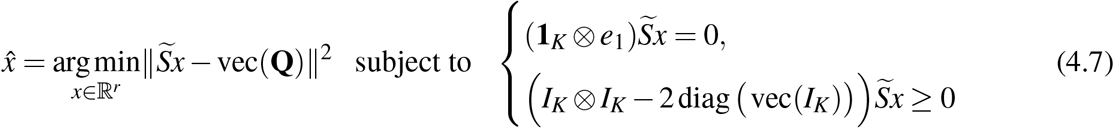

is available from quadratic programming with all linear constraints[46]. Here ⊗ is the Kronecker product, **1**_*K*_ = (1,…, 1) ∈ ℝ^*K*^, *e*_1_ = (1, 0, …, 0) ∈ ℝ^*K*^, diag(·) generates a diagonal matrix, and the last inequality holds entry-wise. The adjusted generator matrix 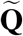 can be obtained by reshaping the vector 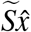 according to (4.6).

Though QP and PA both use the property of shared eigen-structures and incorporate the quadratic programming toolkit, the two methods still differ in many ways. First, the QP algorithm explicitly computes the possibly complex eigendecompositions of **P** and **Q**, while PA avoids doing it by using a different idea of parametrization. Second, the QP algorithm is supposed to find **Q** with eigenvectors that are sufficiently close to those of **P**, while PA forces the eigenvectors of **P** and **Q** to be strictly the same. In addition, due to the different parametrizations, PA also reduces the dimension of the search domain to *O*(*K*), compared to *O*(*K*^2^) in QP.

## V. TOY MODEL

In order to examine the workings of different generator matrix estimators, we use a 1D bistable toy model, whose dynamics is modeled by an overdamped Langevin equation using the Euler-Maruyama[47] method with parameters: temperature *T* = 298*K, mass* = 1, collision frequency *γ* = 1, and time step *δt* = 10^−6^, where time and position are unitless. The potential was of the form

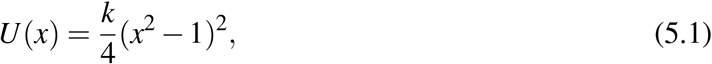

where 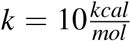. For sampling, 48 sets of simulations were performed. The starting points were equidistantly distributed between *x* = −2 and 2 with increments of Δ*x* = 4*/*47. For each of the 48 starting points we ran 100 repeats of simulations, each for 1,000,000,000 steps. After performing the simulations we created empirical transitions matrices for various lag times, which were then used as the seed information for our algorithms and subsequent detailed balance free energy calculations.

In order to test the efficacy of each algorithm we decided to vary lag time and trajectory lengths. Trajectory length was varied by cutting off our counting statistics for each copy at the appropriate point. For instance, a 10% trajectory length (denoted by *L* = 10%) includes the first tenth of each trajectory. For lag time, we utilized a sliding window approach in creating our empirical transition matrix. In this approach, a transition from state *i* at any time *t* to state *j* at time *t* + *τ* would count towards *N*_*i, j*_. The total number of transition observations based on a given trajectory would be *τ/δt* data points less than the total number of data points in the trajectory. The sliding window approach guarantees a minimal dependence on the lag time for the number of observations to avoid bias.

From the results of our algorithms we calculated the percent error from our expected values, particularly we expected a total Δ*G* of 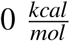 between the two energy minima at *x* = ±1, and a Δ*G* of 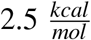 between the energy barrier at *x* = 0 and the average of the two energy minima at *x* = ±1.

## VI. RESULTS AND DISCUSSION

### A. Bistable Potential

We used the data generated for the toy model above to estimate the generator matrix based on various lengths of trajectories and with various lag times using eight algorithms discussed above including DA, WA, QOG, CWO, EM, MLE, QP, and PA. Before we begin it is worth noting the approximate relative computational costs of the algorithms. The DA, and WA algorithms were very low cost, with QOG and CWO only slightly more expensive. EM and MLE were very expensive, requiring specialized computing resources to run for long periods of time. Finally, the QP and PA algorithms were relatively low cost, similar to DA and WA but somewhat more expensive.

The robustness of these methods in terms of their performance was tested by varying lag time and trajectory length. First, we used a fixed trajectory length (in this case, *L* = 10%) to examine the dependence on the lag time. Figure 2A-C shows the estimated free energies along with the analytical potential energy for the toy model. At low *τ*, all algorithms made reasonable predictions, at least in terms of their free energy estimates, with deviations from the model potential function occurring when |*x*| *>* 1.5. At longer lag times, CWO was the first algorithm to break down, which can be seen visibly in the free energy profiles based on *τ* = 50*δt* and greater. The longer lag times increased the CWO free energy estimate for the local minima at *x* = 1. At the longest lag time in panel C, the QP algorithm began to greatly underestimate the extremes of the free energy landscape. Longer lag times tended to make QP less reliable as an estimator for values of |*x*| *>* 1.5.

**FIG. 2.**
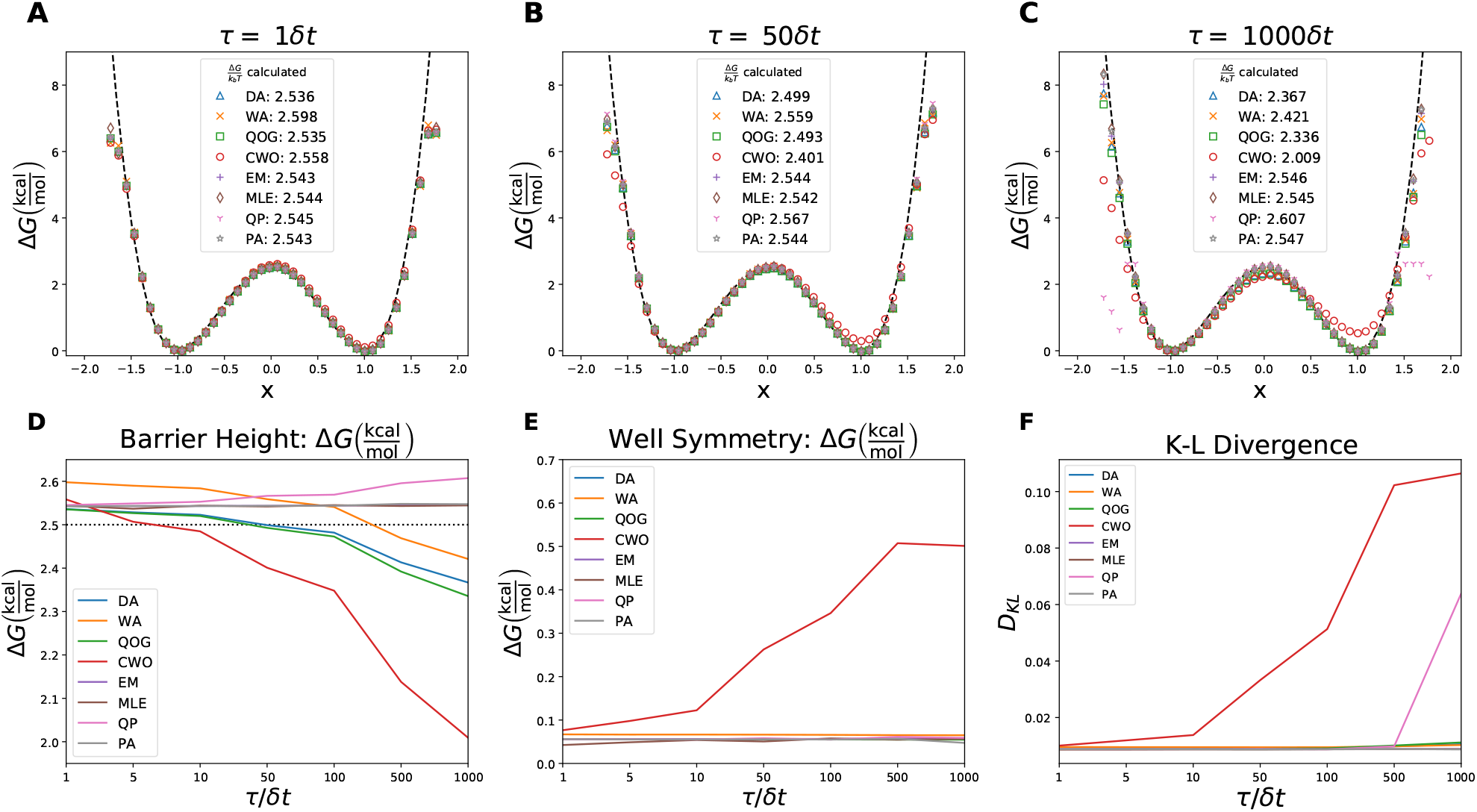
(A-C) Estimated free energy along *x* from DA (blue triangle), WA (orange x), QOG (green square), CWO (red circle), EM (purple +), MLE (brown diamond), QP (pink Y), and PA (gray ★) algorithms for various lag times (expressed in terms of the number of time steps) based on 10% of the data (*L* = 10%); (A) *τ* = *δt*, (B) *τ* = 50*δt*, (C) *τ* = 1000*δt*. The reference black dashed line is the model potential: 2.5(*x*^2^ − 1)^2^. (D) The estimated energy barrier height as a function of *τ*, with an analytical value of 2.5 shown as a black dotted line. (E) The absolute difference of the energy minima as a function of *τ* with an analytical value of 0. (F) The *D*_*KL*_ of the estimated positional probability from the analytic positional probability as a function of *τ*.

Panels D-F in Figure 2 more clearly demonstrate the observed *τ*-dependent behavior of barrier height (defined as 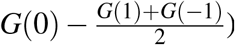) and well symmetry (defined as |*G*(1) − *G*(−1)|) as important features of the free energy profile. All models gave reasonable barrier height estimates at low *τ* with slight over-estimations, WA being the highest. DA, WA, CWO and QOG estimates all decreased with increasing *tau*, where CWO had the fastest rate of decrease, culminating in the worst barrier height estimate of all the algorithms. The EM, MLE, QP, and PA models had slight overestimates of barrier height that remained consistent for all *τ* values, though the QP estimates increased above these other algorithms at the longest lag times. Well symmetry results in Figure 2E, were consistently low for all models except CWO which increased to a plateau with increasing *τ*. The qualitative observation of poor performance in well symmetry for the CWO algorithm was confirmed with Kullback-Leibler divergence[48] (*D*_*KL*_), shown in Figure 2F. *D*_*KL*_ is a measure of difference between probability distributions. The positional probability inside the bistable potential, *U* (*x*), was calculated as

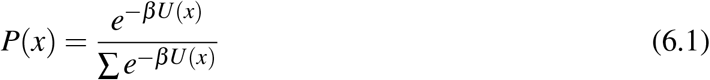

for the analytic potential energy and the estimated potential energy from the different optimization algorithms. The divergence of the estimated *D*_*KL*_ was calculated by the relation

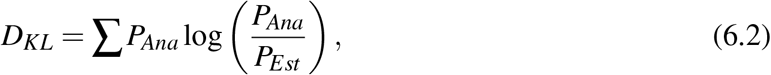

where *P*_*Ana*_ was the analytic probability and *P_Est_* was the estimated probability. *D*_*KL*_ for other values of *τ* are shown in the other panels of Figure S5. The well symmetry and *D*_*KL*_ were qualitatively similar, even though the well symmetry was only using information on the basins. One exception was the deviations in free energy estimates by the QP algorithm at long lag times that are captured in the *D*_*KL*_ figure but not the well symmetry.

Figure 3A-C illustrates the free energy profiles estimated from various methods for a fixed *τ* (*τ* = 10*δt*) and varying trajectory length. Using all of the trajectory, all the algorithms provided reasonable estimates for the free energy. Deviations in the basin of the well at *x* = 1 appeared as the length of the trajectory was reduced, as well as poor estimations at the extremes of the potential, |*x*| *>* 1.5.

**FIG. 3.**
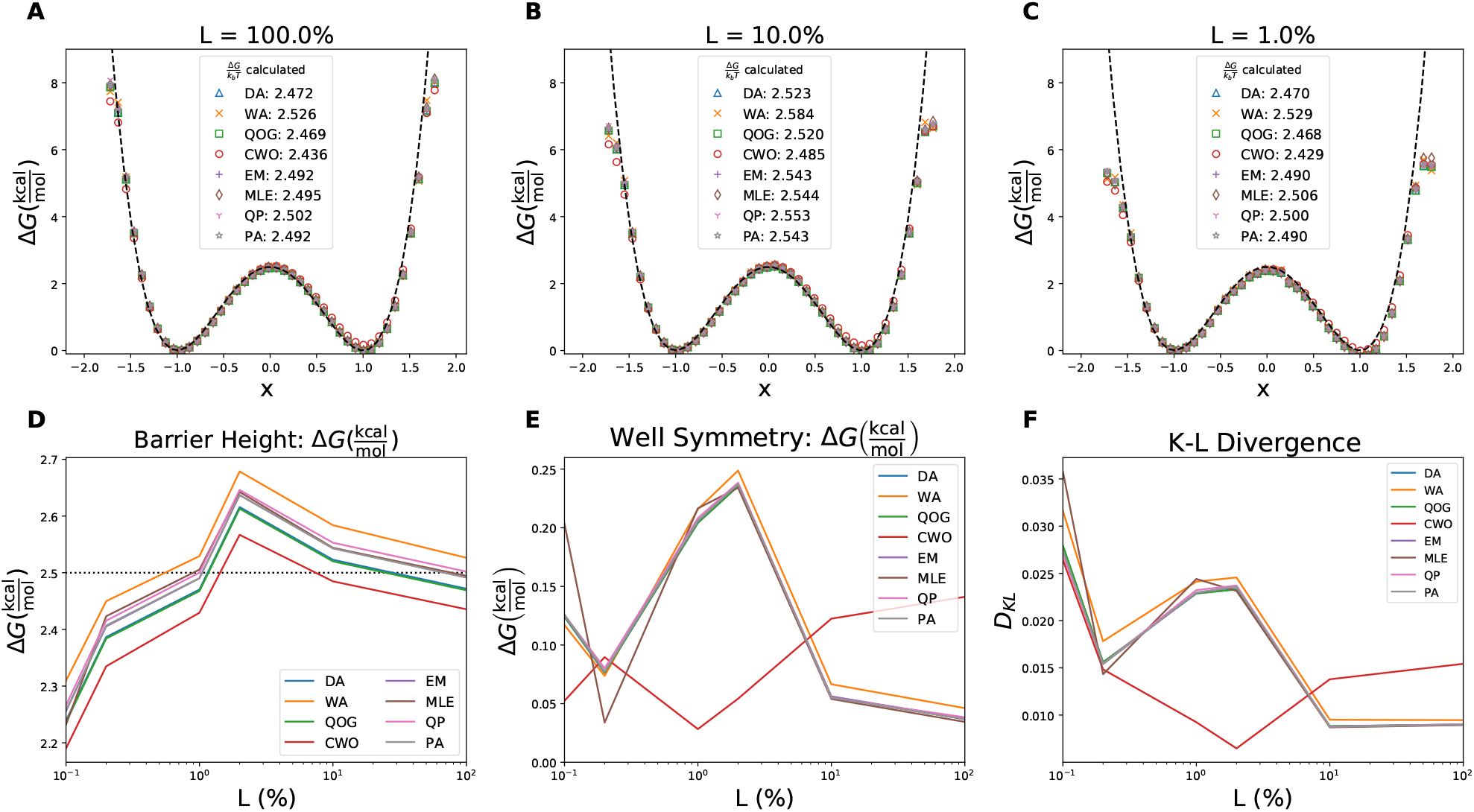
(A-C) Estimated free energy along *x* from DA (blue triangle), WA (orange x), QOG (green square), CWO (red circle), EM (purple +), MLE (brown diamond), QP (pink Y), and PA (gray *\**) for various trajectory lengths; (A) *L* = 100%, (B) *L* = 10%, (C) *L* = 0.1%. (D) The energy barrier height Δ*G* (as a function of trajectory length) with an analytical value of 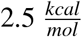, shown as a black dotted line for reference. (E) The absolute difference of the energy minima as a function of *L*% with an analytical value of 0. (F) *D*_*KL*_ of the estimated positional probability from the analytic positional probability as a function of trajectory length.

All algorithms had qualitative similarities in their estimates of barrier height in Figure 3D, with low estimates for the shortest trajectory increasing the a peak at *L* = 5%, then approaching the true value at *L* = 100. The relative spacing of the barrier height remained mostly consistent for a given trajectory length. The most accurate algorithms for the full trajectory were EM, MLE, QP, and PA. WA overestimated the barrier height, though this behavior was dependent on the *τ* chosen, while DA, QOG, and CWO algorithms all underestimated the barrier height. At the lowest trajectory length, *L* = 0.1%, the estimations of barrier peak was inaccurate, all significantly below the analytic value of 2.5. Differences between the related QP and PA algorithms were also observed at lower trajectory lengths, with QP giving slightly higher estimates than PA.

When we used shorter trajectories, the barrier estimates deviated from the correct values. From Figure 3D, the estimated barriers were flatter for the shortest trajectories, *L <* 1%. This can be easily understood from observing that with shorter trajectory lengths, there was more dependence on the starting point of trajectories, which were distributed evenly over the space. As the trajectory length shortens, the free energy profiles resemble more the initial flat distribution. A similar overestimation of the barrier at *L* = 2% for all algorithms indicated further the dependence on the data for accurate results.

The well symmetry and *D*_*KL*_ panels in Figure 3E and F confirmed earlier observations that CWO behaved differently than the other algorithms. The *D*_*KL*_ values in Figure 3F confirmed that CWO performed poorly at the longer trajectory lengths, but at *L <* 2% the CWO deviations was comparable or superior to the other algorithms. *D*_*KL*_ for other trajectory lengths are shown in the other panels of Figure S5.

Tables S1 and S2 provide more extensive data on the dependence of these algorithms on lag time and trajectory length as they summarize various estimates based on varying both quantities. In the tables one sees at the first row for each method, the errors associated with the barrier height and well symmetry estimates are listed in Tables S1 and S2, respectively for full-length trajectories (*L* = 100%) but varying lag times. In all cases, the barrier height errors were quite small at *τ* = *δt* but the well symmetry error was considerably higher than the rest for CWO (by a factor of 17) at *L* = 100% and *τ* = *δt*. At the *L* = 0.01% and *τ* = 1000*δt*, CWO estimates of well symmetry improved relative to the other algorithms. The well symmetry errors were generally negligible in most cases except for CWO methods. However, barrier height estimates worsen in most cases as the trajectory length decreases or as lag time increases. At *L* = 0.1%, all algorithms perform similarly in barrier height estimates.

The above analysis implies that most algorithms above (except for CWO) were robust for estimating thermodynamic quantities (here the free energy difference between the minima) than kinetic ones (here the barrier height, which is related to activation energy in chemical kinetics literature). We also examined the performance of these methods in terms of estimating MFPT associated with the transition from the energy minimum at *x* = −1 to the energy minimum at *x* = 1 (i.e., from the left to the right well). The analytical MFPT is 34.3. This value is estimated from Relation (2.4), using *G*(*x*) = *U* (*x*) (from Relations (5.1) and *D* = *k*_*B*_*T* = 0.59. To estimate the MFPT from the simulations, we employ Relations (2.5), which requires an estimate for the free energies (*G*_*i*_) as well as *D*_*i*_. Table S3 provides an estimate for *D*(*x*) (averaged over *x*) from generator matrices that are estimated by DA, WA, QOG, CWO, EM, MLE, QP, and PA. Generally, the estimates were influenced more by *τ* than the trajectory lengths. Unlike the free energy estimates that tend to be more accurate for lower *τ* values, the diffusion constant estimates were more accurate for longer lag times. The DA and QOG algorithms generated the closest estimates to the analytical diffusion, using long lag times, though they still overestimate by approximately 80%. The QP and PA algorithms were the least accurate estimators at the long lag times, but all algorithms gave results within the same order of magnitude.

MFPT estimates based on *D*_*i*_ and *G*_*i*_ estimates are shown in Table I for varying lag times and trajectory lengths. As compared to the analytical value (34.4), all estimates were somewhat lower than expected, which is the direct result of the overestimation of *D*(*x*). Similar to *D*(*x*) and contrary to *G*(*x*), the estimates were more accurate at longer lag times. The MFPT estimates were also affected by the trajectory length. For *L* = 100% and *L* = 10%, the difference was not significant between the MFPT estimates; however, at *L* = 1% most methods start to show more deviation from the actual MFPT values as compared to *L* = 10% and *L* = 100% estimates. The algorithms with the closest MFPT estimates were DA, WA, and QOG, but even the closest values were still half of the analytic MFPT. Similar to diffusion, QP and PA had the least accurate MFTP estimates, but they were still within a reasonable range compared to the other algorithms.

The generator matrix contains both thermodynamic and kinetic information as the above analysis clearly illustrates. The eight different generator matrix estimators that we used here had their own pros and cons. The CWO method was a relatively poor barrier height estimator as compared to the other methods used here, especially at long lag times. On the other hand, it’s well symmetry estimation surpassed other algorithms when the trajectory length was short. Furthermore, CWO performed similarly to other methods for estimating diffusion constant and MFTP. The EM, MLE, and PA algorithms was generally robust for estimating both thermodynamic and kinetic properties, with accurate predictions of barrier height for all lag times using the full trajectory. Additionally, the diffusion and MFPT estimates for EM, MLE, and PA were in general agreement with the other algorithms. It is important to remember that the PA algorithm was able to achieve these results while requiring significantly less computing time.

**TABLE I.**
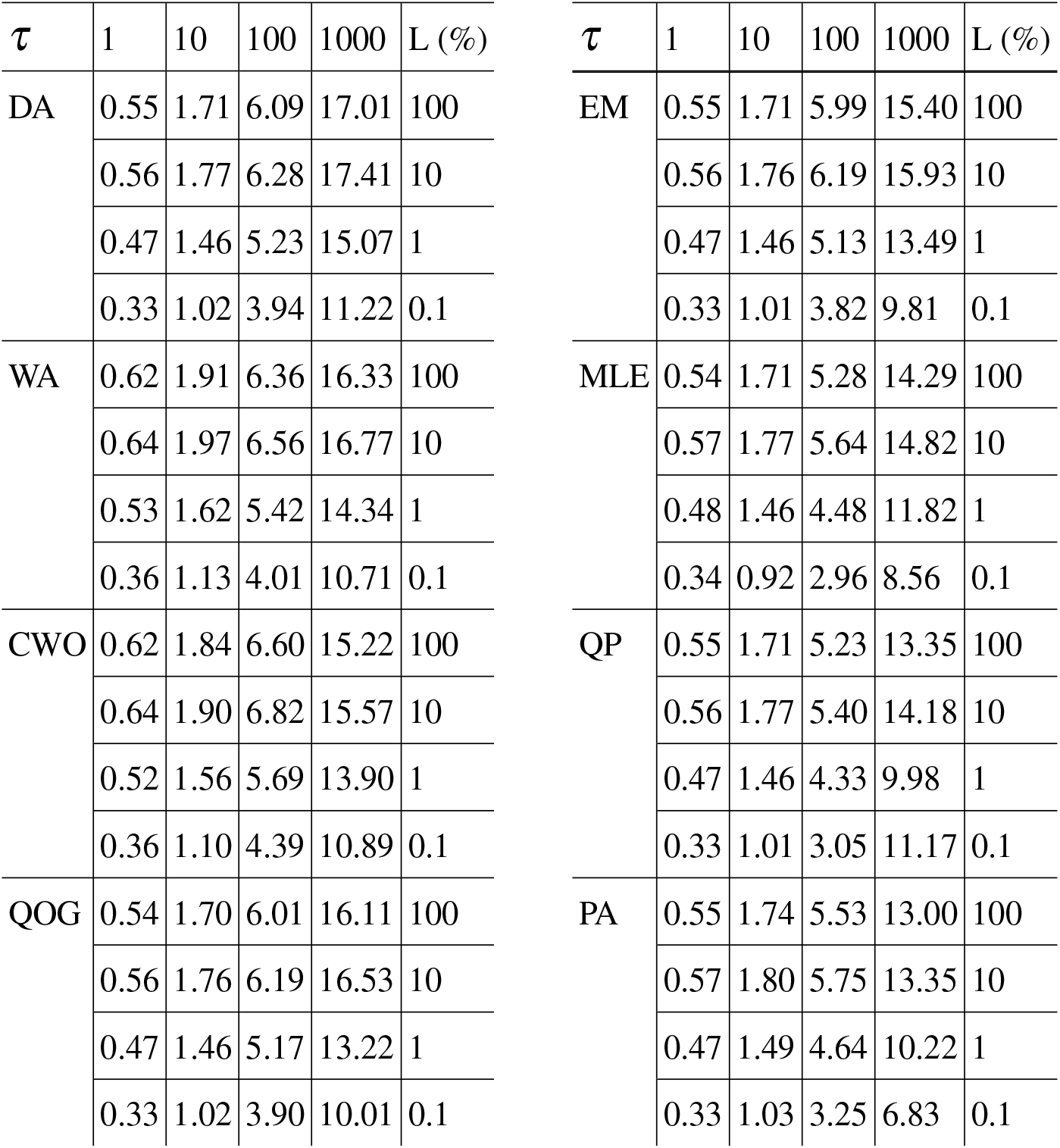
MFPT predicted using various estimates of the generator matrix including: DA, WA, QOG, CWO, EM, MLE, QP, and PA. Increasing values of *τ* are seen along the columns while decreasing trajectory length (*L*) are represented along the rows. The analytical MFPT is 34.3.

In these results, there were certain features that were broadly true. For instance, more data lead to better estimates; although different algorithms had different sensitivity levels to the amount of data used. An interesting observation is that free energy estimates were generally more accurate for shorter lag times but the diffusion constant estimates were more accurate for longer lag times. The overestimation of diffusion constant (and subsequent underestimation of the MFPT) can be attributed to the correlatedness of the data generated by the overdamped langevin equation. In a typical molecular dynamics simulations, we also expect to observe such correlated behavior, which would result in similar correlatedness in the data. The longer lag times reduced correlation in the empirical transition matrix. On the other hand, longer lag times result in empirical transition matrices that have more non-zero off-diagonal elements and are farther away from the generator matrix, which in this case is expected to be tridiagonal.

## VII. CONCLUSIONS

We implemented eight algorithms for overcoming the “embeddability problem” in MSMs. The efficiency of these algorithms were tested on a 1D bistable toy model. The relative performance of each algorithm were compared with regards to their ability to handle differing lag times and trajectory lengths. We found that depending on whether one was interested in free energy calculations or kinetic characterization of a process, different algorithms may behave differently. For thermodynamic characterization of a process, EM, MLE, and PA showed robust results for different lag times. PA was able to achieve comparable results in kinetic information estimation to EM and MLE despite having a significantly lower computational cost. CWO performed the worst when all the data was used, though it may perform better when trajectories are short and lag times are long.

In terms of kinetic estimations such as MFPT and diffusion constant, the simplest algorithm, DA, gave marginally closer estimates, though all algorithms performed similarly. In general, in the abundance of data, all algorithms provide reasonable results; however, the lag time dependence should be checked by repeating the analysis for various lag times to ensure the reliability of the results.

The development of the novel PA algorithm, while conceptually similar to the QP algorithm, improved upon the long lag time errors of QP in free energy estimation. PA also had similarly accurate estimates for barrier height and well symmetry to more computationally expensive algorithms like EM and MLE. This represents a viable approach to extracting embedded information for large transition matrices when computational resources are at a premium.

Overall, this exploration provides the interested reader a foundation for dealing with kinetic and thermodynamic analysis of equilibrium MD trajectories, particularly in the context of MSM. Through our analysis we hope to shed some light on the approaches as well as the information which can be garnered from direct analysis of MD data. More data is almost always preferable but in situations where this is not possible, overcoming the embeddability problem is a possibility for estimating the thermodynamic and kinetic quantities reliably.

## Data Availability

The data that support the findings of this study are available from the corresponding author upon reasonable request.

## Acknowledgement

This research is supported by the National Science Foundation under Awards 1940188, 1945465, 1934985, 1940124, 1940276, and 1940223. This research is also supported by the Arkansas High Performance Computing Center which is funded through multiple National Science Foundation grants and the Arkansas Economic Development Commission.

