## Supporting Information for "Addressing the Embeddability Problem in Transition Rate Estimation"

### Supplemental Information

#### BARRIER HEIGHT

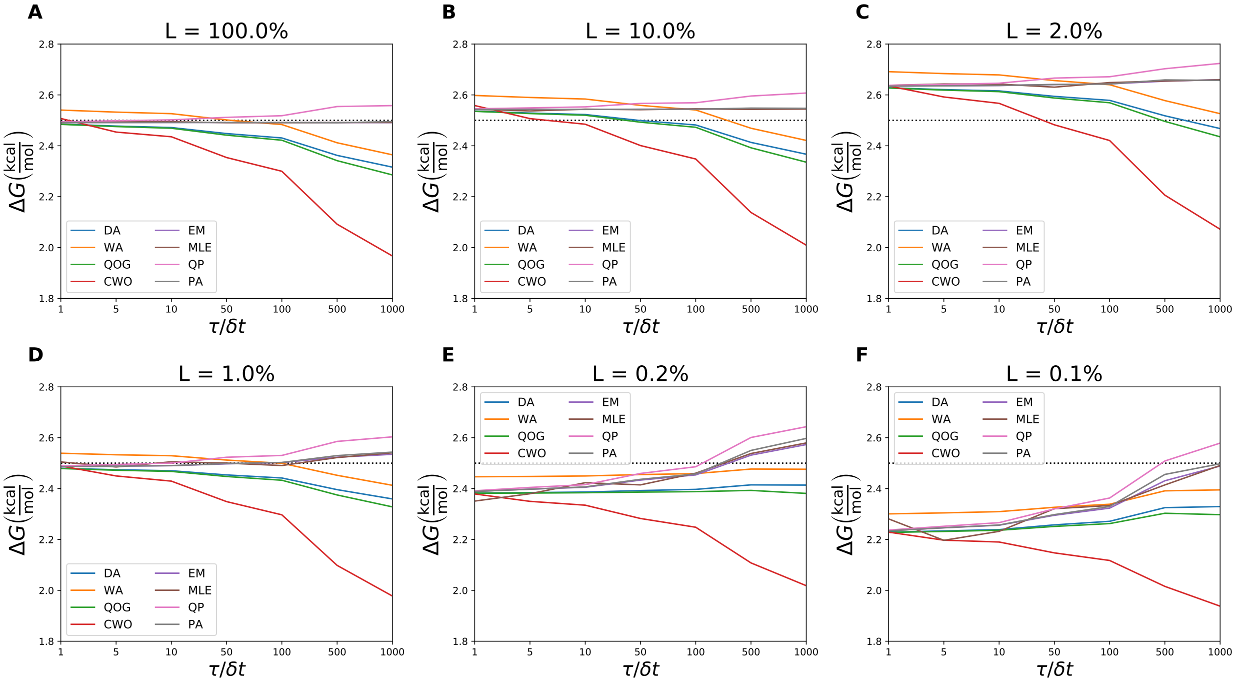


FIG. S1. (A-E) Barrier height as a function of $\tau/\delta t$ with the trajectory length, $L\%$, held at a different values in each plot; (A) $L = 100\%$, (B) $L = 10\%$, (C) $L = 2\%$, (D) $L = 1\%$, (E) $L = 0.2\%$, (F) $L = 0.1\%$. The algorithms shown are DA, PA, QOG, CWO, EM, MLE, QP, and PA.

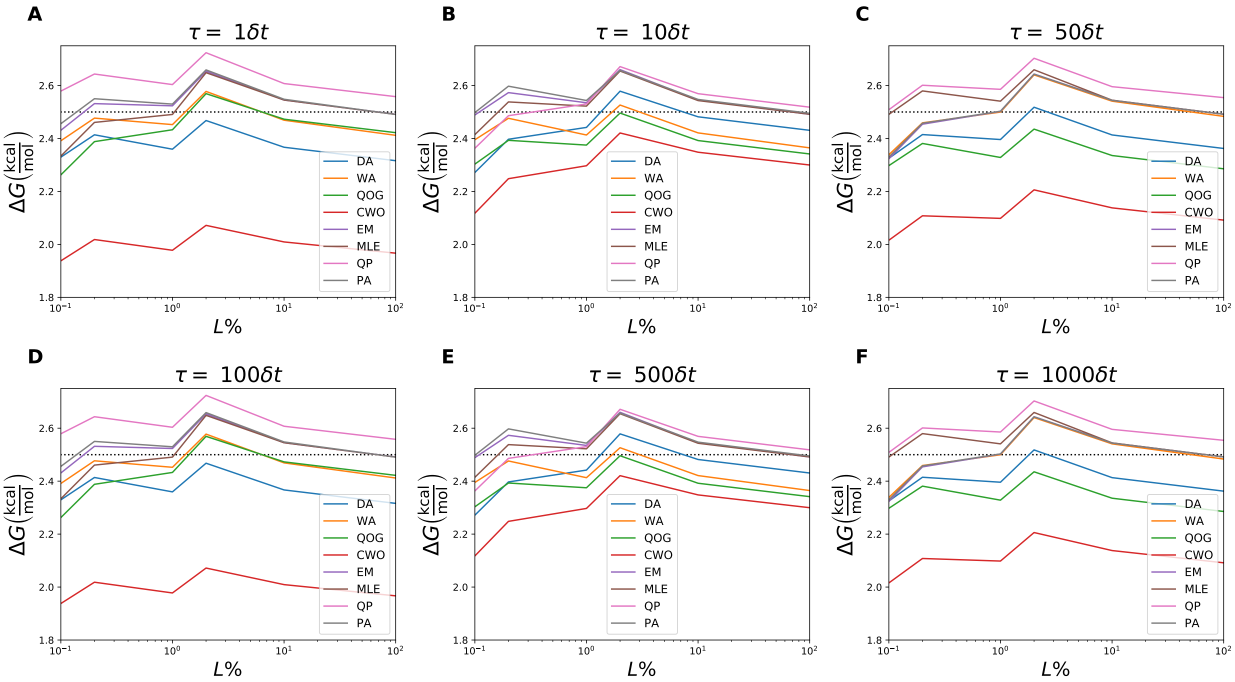


FIG. S2. (A-E) Barrier height as a function of simulation length, plotted on a logarithmic scale, with $\tau$ held at a different value in each plot; (A) $\tau = \delta t$, (B) $\tau = 10\delta t$, (C) $\tau = 50\delta t$, (D) $\tau = 100\delta t$, (E) $\tau = 500\delta t$, (F) $\tau = 1000\delta t$. The algorithms shown are DA, PA, QOG, CWO, EM, MLE, QP, and PA.

#### WELL SYMMETRY

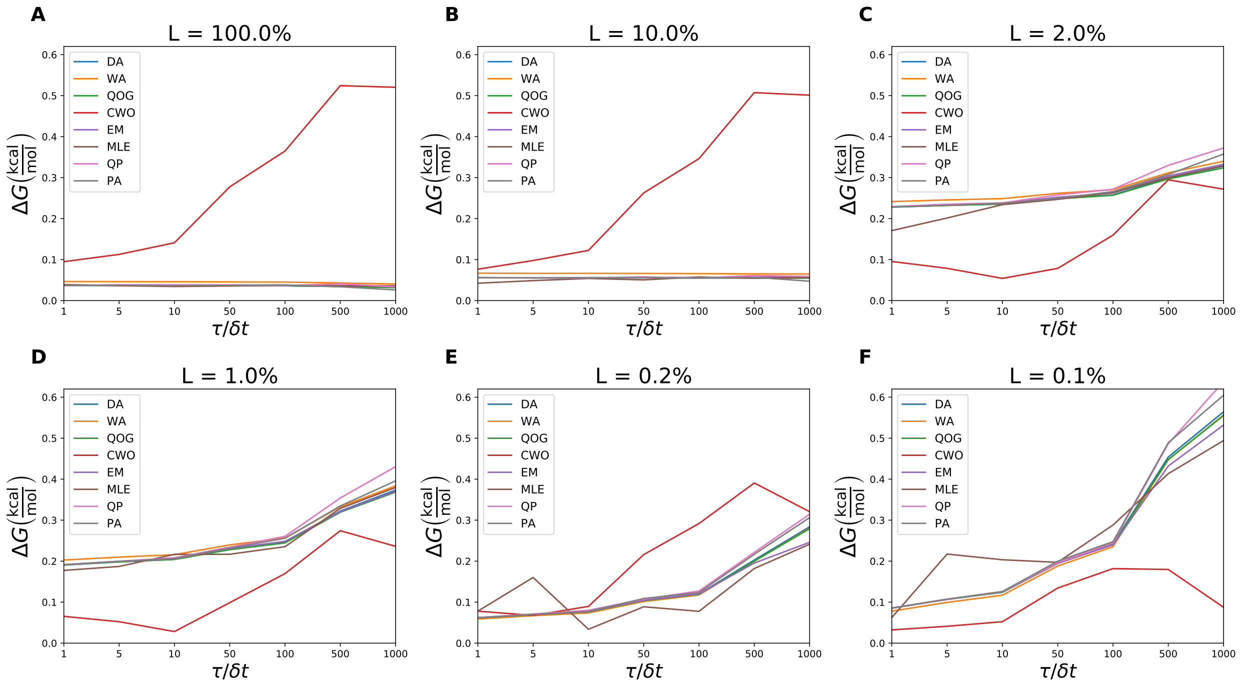


FIG. S3. A-E) Well symmetry as a function of $\tau/\delta t$ with the trajectory length, $L\%$, held at a different values in each plot; (A) $L = 100\%$, (B) $L = 10\%$, (C) $L = 2\%$, (D) $L = 1\%$, (E) $L = 0.2\%$, (F) $L = 0.1\%$. The algorithms shown are DA, PA, QOG, CWO, EM, MLE, QP, and PA.

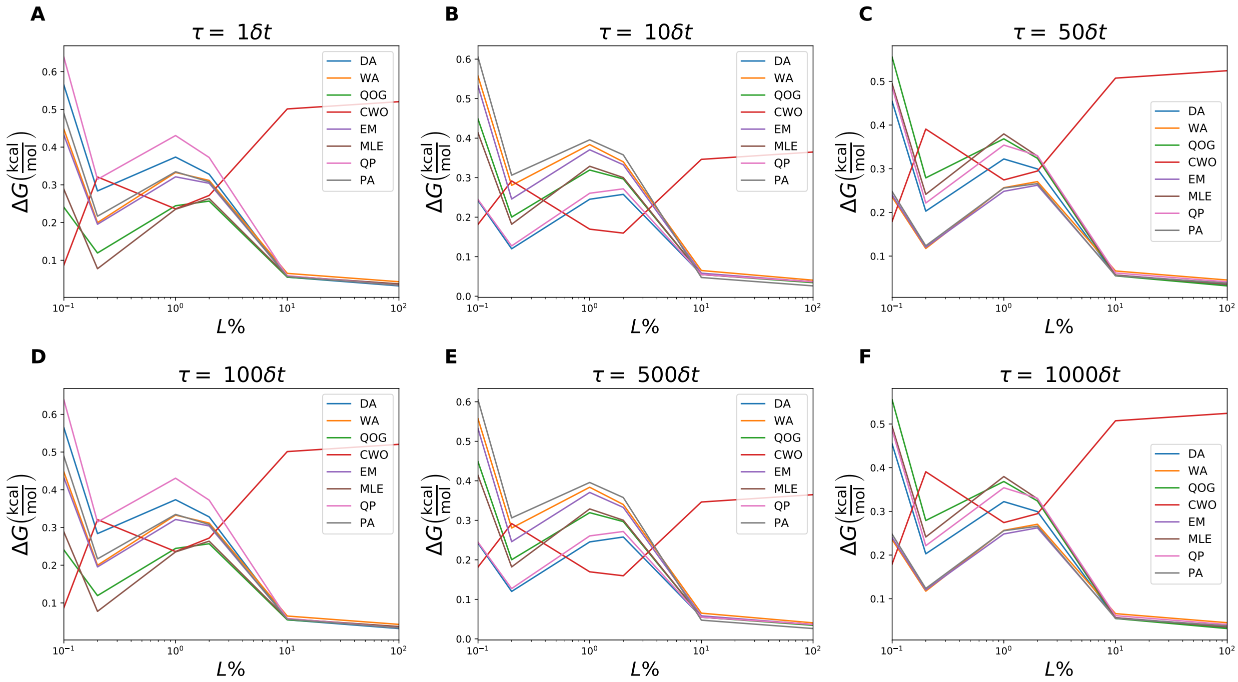


FIG. S4. (A-E) Well symmetry as a function of simulation length, plotted on a logarithmic scale, with $\tau$ held at a different value in each plot; (A) $\tau = \delta t$, (B) $\tau = 10\delta t$, (C) $\tau = 50\delta t$, (D) $\tau = 100\delta t$, (E) $\tau = 500\delta t$, (F) $\tau = 1000\delta t$. The algorithms shown are DA, PA, QOG, CWO, EM, MLE, QP, and PA.

#### KULLBACK-LEIBLER DIVERGENCE

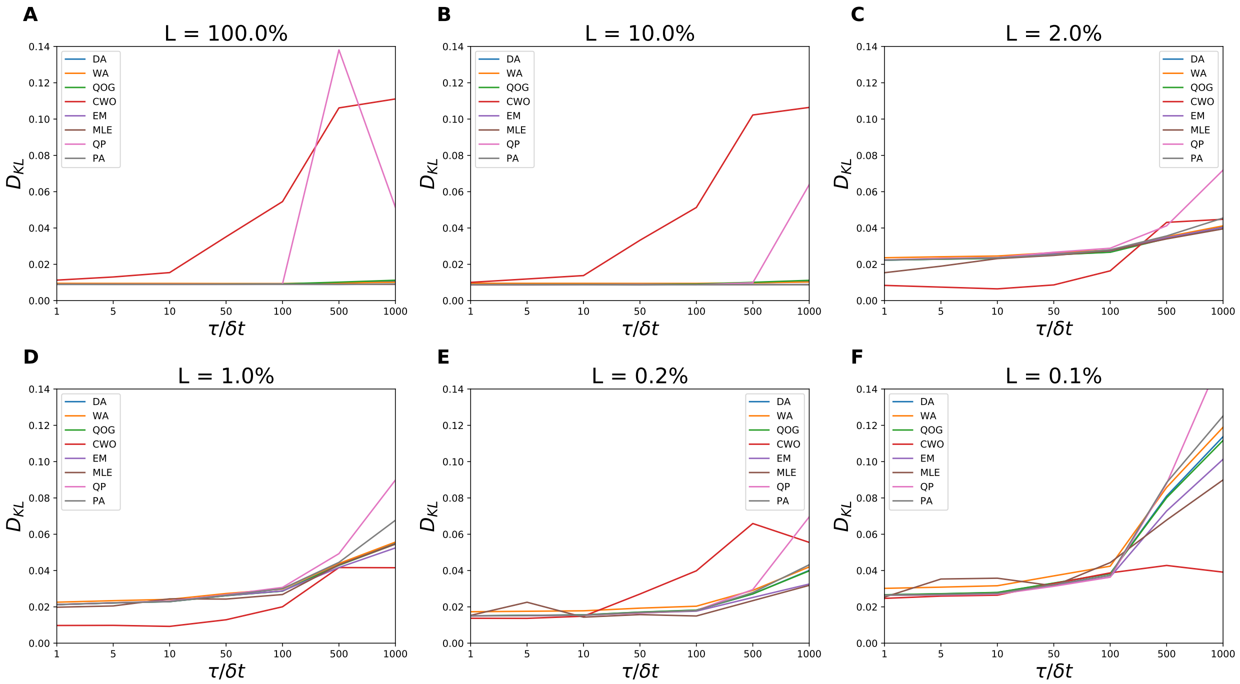


FIG. S5. (A-E) $D_{KL}$ as a function of $\tau/\delta t$ with the trajectory length, $L\%$, held at a different values in each plot; (A) $L = 100\%$, (B) $L = 10\%$, (C) $L = 2\%$, (D) $L = 1\%$, (E) $L = 0.2\%$, (F) $L = 0.1\%$. The algorithms shown are DA, PA, QOG, CWO, EM, MLE, QP, and PA.

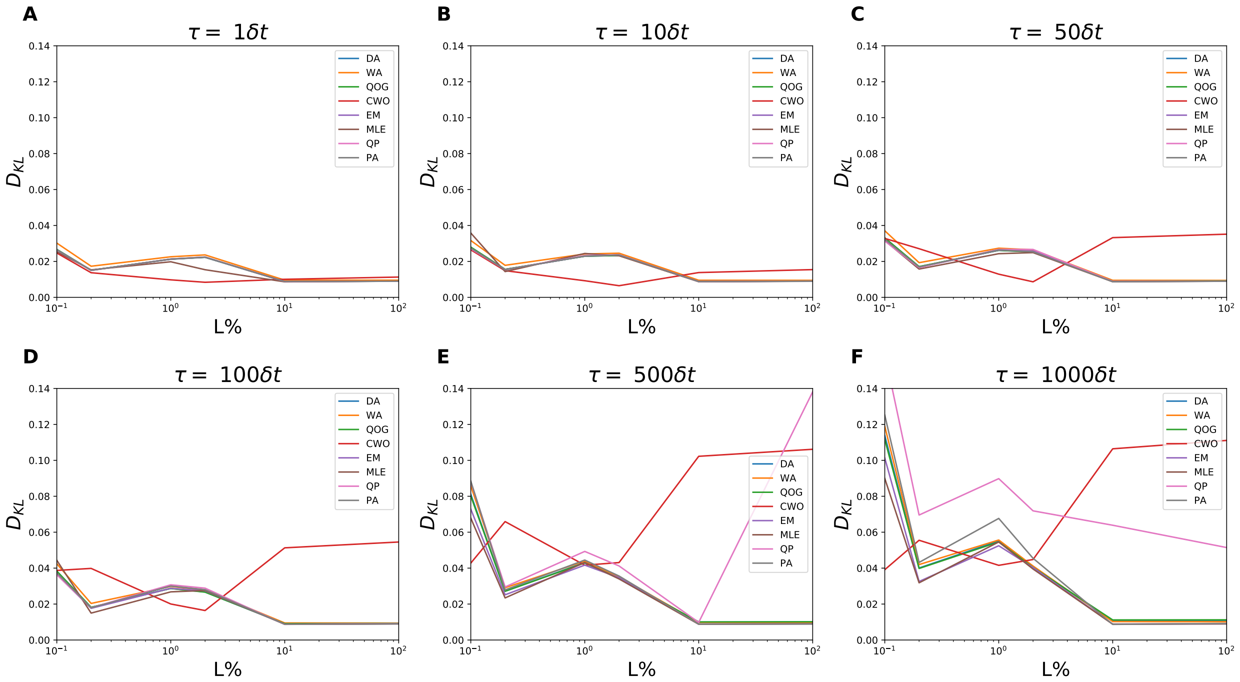


FIG. S6. (A-E) $D_{KL}$ as a function of simulation length, plotted on a logarithmic scale, with $\tau$ held at a different value in each plot; (A) $\tau = \delta t$, (B) $\tau = 10\delta t$, (C) $\tau = 50\delta t$, (D) $\tau = 100\delta t$, (E) $\tau = 500\delta t$, (F) $\tau = 1000\delta t$. The algorithms shown are DA, PA, QOG, CWO, EM, MLE, QP, and PA.

#### DATA TABLES

TABLE S1. Percent errors in barrier height ($\Delta G = G(0) - \frac{G(-1)+G(1)}{2}$) based on different free energy predictions. The predictions are based on various estimates of the generator matrix including: DA, WA, QOG, CWO, EM, MLE, QP, and PA. Increasing values of $\tau$ are seen along the columns while decreasing trajectory length ($L$) are represented along the rows.

| $\tau$ | 1 | 10 | 100 | 1000 | L (%) |
|---|---|---|---|---|---|
| DA | 0.59 | 1.13 | 2.77 | 7.36 | 100 |
| | 1.45 | 0.92 | 0.73 | 5.33 | 10 |
| | 0.81 | 1.19 | 2.33 | 5.62 | 1 |
| | 10.84 | 10.45 | 9.14 | 6.82 | 0.1 |
| WA | 1.62 | 1.05 | 0.66 | 5.42 | 100 |
| | 3.92 | 3.35 | 1.63 | 3.16 | 10 |
| | 1.56 | 1.17 | 0.01 | 3.48 | 1 |
| | 7.97 | 7.62 | 6.46 | 4.20 | 0.1 |
| CWO | 0.29 | 2.58 | 8.01 | 21.33 | 100 |
| | 2.33 | 0.61 | 6.08 | 19.63 | 10 |
| | 0.44 | 2.82 | 8.12 | 20.89 | 1 |
| | 10.84 | 12.40 | 15.30 | 22.49 | 0.1 |
| QOG | 0.63 | 1.24 | 3.12 | 8.58 | 100 |
| | 1.42 | 0.80 | 1.09 | 6.58 | 10 |
| | 0.84 | 1.29 | 2.68 | 6.87 | 1 |
| | 10.89 | 10.55 | 9.50 | 8.10 | 0.1 |

| $\tau$ | 1 | 10 | 100 | 1000 | L (%) |
|---|---|---|---|---|---|
| EM | 0.34 | 0.33 | 0.34 | 0.36 | 100 |
| | 1.72 | 1.73 | 1.76 | 1.83 | 10 |
| | 0.56 | 0.39 | 0.10 | 1.40 | 1 |
| | 10.63 | 9.76 | 7.07 | 0.45 | 0.1 |
| MLE | 0.36 | 0.22 | 0.36 | 0.34 | 100 |
| | 1.75 | 1.76 | 1.81 | 1.78 | 10 |
| | 0.20 | 0.22 | 0.37 | 1.63 | 1 |
| | 8.76 | 10.72 | 6.66 | 0.36 | 0.1 |
| QP | 0.23 | 0.08 | 0.73 | 2.32 | 100 |
| | 1.81 | 2.12 | 2.77 | 4.29 | 10 |
| | 0.43 | 0.01 | 1.22 | 4.14 | 1 |
| | 10.52 | 9.35 | 5.48 | 3.16 | 0.1 |
| PA | 0.34 | 0.34 | 0.34 | 0.20 | 100 |
| | 1.70 | 1.72 | 1.75 | 1.89 | 10 |
| | 0.55 | 0.38 | 0.11 | 1.74 | 1 |
| | 10.64 | 9.73 | 6.82 | 0.09 | 0.1 |

TABLE S2. Absolute errors in $\Delta G = |G(1) - G(-1)|$, the free energy difference between the two minima, with an analytical value of 0. The predictions are based on various estimates of the generator matrix including: DA, WA, QOG, CWO, EM, MLE, QP, and PA. Increasing values of $\tau$ are seen along the columns while decreasing trajectory length ($L$) are represented along the rows.

| $\tau$ | 1 | 10 | 100 | 1000 | L (%) |
|---|---|---|---|---|---|
| DA | 0.04 | 0.04 | 0.04 | 0.03 | 100 |
| | 0.06 | 0.06 | 0.06 | 0.06 | 10 |
| | 0.19 | 0.20 | 0.25 | 0.37 | 1 |
| | 0.09 | 0.12 | 0.24 | 0.56 | 0.1 |
| WA | 0.05 | 0.05 | 0.05 | 0.04 | 100 |
| | 0.07 | 0.07 | 0.07 | 0.07 | 10 |
| | 0.20 | 0.22 | 0.26 | 0.38 | 1 |
| | 0.08 | 0.12 | 0.24 | 0.56 | 0.1 |
| CWO | 0.09 | 0.14 | 0.36 | 0.52 | 100 |
| | 0.08 | 0.12 | 0.35 | 0.50 | 10 |
| | 0.07 | 0.03 | 0.17 | 0.24 | 1 |
| | 0.03 | 0.05 | 0.18 | 0.09 | 0.1 |
| QOG | 0.04 | 0.04 | 0.04 | 0.03 | 100 |
| | 0.06 | 0.06 | 0.06 | 0.05 | 10 |
| | 0.19 | 0.20 | 0.24 | 0.37 | 1 |
| | 0.09 | 0.12 | 0.24 | 0.56 | 0.1 |

| $\tau$ | 1 | 10 | 100 | 1000 | L (%) |
|---|---|---|---|---|---|
| EM | 0.04 | 0.04 | 0.04 | 0.04 | 100 |
| | 0.06 | 0.06 | 0.06 | 0.06 | 10 |
| | 0.19 | 0.21 | 0.25 | 0.37 | 1 |
| | 0.09 | 0.13 | 0.24 | 0.53 | 0.1 |
| MLE | 0.04 | 0.03 | 0.04 | 0.04 | 100 |
| | 0.04 | 0.05 | 0.06 | 0.06 | 10 |
| | 0.18 | 0.22 | 0.24 | 0.38 | 1 |
| | 0.06 | 0.20 | 0.29 | 0.49 | 0.1 |
| QP | 0.04 | 0.04 | 0.04 | 0.03 | 100 |
| | 0.06 | 0.06 | 0.05 | 0.06 | 10 |
| | 0.19 | 0.21 | 0.26 | 0.43 | 1 |
| | 0.09 | 0.13 | 0.24 | 0.64 | 0.1 |
| PA | 0.04 | 0.04 | 0.04 | 0.03 | 100 |
| | 0.06 | 0.06 | 0.06 | 0.05 | 10 |
| | 0.19 | 0.21 | 0.26 | 0.40 | 1 |
| | 0.09 | 0.13 | 0.25 | 0.60 | 0.1 |

TABLE S3. Diffusion constant $D(x)$ averaged over $x = -2$ to $x = 2$ and estimated from $D_i$ values. Increasing values of $\tau$ are seen along the columns while decreasing trajectory length ($L$) are represented along the rows. The predictions are based on various estimates of the generator matrix including: DA, WA, QOG, CWO, EM, MLE, QP, and PA. The underlying model has an analytical value of $D(x) = 0.59$ for all $x$.

| $\tau$ | 1 | 10 | 100 | 1000 | L (%) |
|---|---|---|---|---|---|
| DA | 36.54 | 11.49 | 3.57 | 1.06 | 100 |
| | 36.69 | 11.54 | 3.59 | 1.06 | 10 |
| | 36.65 | 11.57 | 3.60 | 1.06 | 1 |
| | 35.71 | 11.31 | 3.51 | 1.06 | 0.1 |
| WA | 36.81 | 11.68 | 3.73 | 1.21 | 100 |
| | 36.96 | 11.72 | 3.74 | 1.21 | 10 |
| | 36.91 | 11.75 | 3.75 | 1.20 | 1 |
| | 35.92 | 11.47 | 3.66 | 1.21 | 0.1 |
| CWO | 36.72 | 11.70 | 3.78 | 1.22 | 100 |
| | 36.86 | 11.75 | 3.80 | 1.22 | 10 |
| | 36.82 | 11.78 | 3.81 | 1.22 | 1 |
| | 35.85 | 11.52 | 3.71 | 1.22 | 0.1 |
| QOG | 36.56 | 11.51 | 3.59 | 1.08 | 100 |
| | 36.70 | 11.55 | 3.60 | 1.08 | 10 |
| | 36.66 | 11.58 | 3.61 | 1.07 | 1 |
| | 35.72 | 11.32 | 3.52 | 1.08 | 0.1 |

| $\tau$ | 1 | 10 | 100 | 1000 | L (%) |
|---|---|---|---|---|---|
| EM | 36.83 | 11.78 | 3.86 | 1.38 | 100 |
| | 36.97 | 11.83 | 3.88 | 1.38 | 10 |
| | 36.94 | 11.86 | 3.89 | 1.38 | 1 |
| | 35.98 | 11.59 | 3.79 | 1.39 | 0.1 |
| MLE | 36.83 | 11.78 | 3.86 | 1.38 | 100 |
| | 36.98 | 11.83 | 3.88 | 1.38 | 10 |
| | 36.86 | 11.85 | 3.89 | 1.38 | 1 |
| | 36.20 | 11.61 | 3.80 | 1.39 | 0.1 |
| QP | 36.94 | 11.90 | 4.02 | 1.82 | 100 |
| | 37.10 | 11.97 | 4.05 | 1.81 | 10 |
| | 37.06 | 12.00 | 4.08 | 1.81 | 1 |
| | 36.11 | 11.72 | 3.97 | 1.87 | 0.1 |
| PA | 36.65 | 11.56 | 3.65 | 1.53 | 100 |
| | 36.78 | 11.60 | 3.65 | 1.54 | 10 |
| | 36.73 | 11.61 | 3.65 | 1.54 | 1 |
| | 35.78 | 11.36 | 3.57 | 1.54 | 0.1 |
